# Writing does not Impact the Evolution of Syntax

**DOI:** 10.1101/2025.07.19.665695

**Authors:** Carlo Y. Meloni, Jessica K. Ivani, Taras Zakharko, Guanghao You, Chundra A. Cathcart, Balthasar Bickel

## Abstract

The acquisition of writing has a lasting effect on how languages are processed in the brain, raising the possibility that it also affects how they are used and evolve over time. A popular hypothesis is that writing specifically leads to increased use and evolution of complex hierarchical structures in language. While most case studies on individual languages have supported this, it remains unclear whether results scale to larger and more diverse sets of languages. Here we test the prediction for language use by modeling hierarchy depth and sentence length in Universal Dependencies corpora from 30 languages, and the prediction for language evolution by modeling the phylogenetic dynamics of structural asymmetry in clause combinations of 59 languages from three families. We found no effect of writing in either analysis. Hierarchy depth and sentence length do not demonstrably differ between spoken and written use of language, and the introduction of writing had no discernible effect on the evolution of clause combining in grammar. There is weak evidence for a decrease in variance after the introduction of writing in one family, possibly reflecting the impact of increased normativity. In conclusion, our findings challenge popular beliefs about hierarchy and suggest that the evolution of syntax is more driven by culturally variable ideas about style in linguistic expression than by the evolution of writing.

---

Learning how to read and write fundamentally restructures language processing, enhancing neural activity in various brain areas [1], reducing the cognitive load imposed by complex syntactic structures [2], improving detection of grammatical norm violations [3], and facilitating prediction even in spoken language comprehension [4]. More broadly, the acquisition of reading and writing is proposed to augment both linguistic and non-linguistic cognitive abilities [5]. Furthermore, unlike spoken communication, writing often lacks shared situational knowledge, non-linguistic context, and interactive cues, necessitating more explicit encoding. Also, writing allows for greater revision and refinement, reducing working memory constraints and allowing more complex structures of expression [6].

It is plausible that these far-reaching neuro-cognitive and behavioral effects would leave an imprint on how language is used and on how it evolves. Changes in the processing system may induce changes in how sentences are planned and executed, promoting syntactic complexity [7], in particular an increase of hierarchical over concatenative syntax [8]. While such effects might first be limited to literate individuals, it is plausible that these would act as socially prominent innovators, facilitating the adoption of more complex syntactic patterns more widely [9] and, as a result, impact how grammars evolved over time.

Potential evidence for effects on use comes from corpus studies of English, suggesting substantial differences between written and oral styles [10, 11, 12]. Consistent with this, studies of oral languages such as Inuktitut [13] and Pirahã [14, 15] suggest that they under-use hierarchical sentence structure (subordination) compared to languages with established writing traditions [16, 17, 18], although other oral languages, for example some languages from Papua New Guinea or Nepal, make heavy use of complex hierarchical structures [19, 20], and the grammars of complex sentences do not seem to cluster along a difference between languages in literate and illiterate societies [21].

It has also been frequently claimed that before the development of writing systems, syntax was predominantly flat, favoring simpler constructions reliant on coordinating (e.g. ‘and’) rather than subordinating (e.g. ‘when’) conjunctions [22, 23]. In contrast, written language has been claimed to exhibit more hierarchical syntax, characterized by complex sentence structures and extensive use of subordinate clauses [24, 25, 26, 8]. Consistent with this, a gradual increase in hierarchy has been documented in several languages, including Akkadian [27], Biblical Hebrew [28, 29], Old English [30, 31], Dutch [32], German [32], and Old Chinese [33]. Taken together, the current evidence rests on individual case studies from a relatively small set of languages. While most studies suggest effects of writing on the use and evolution of hierarchical expression, there are exceptions. Therefore, the generality of effects remains an open question.

Here we put these effects to a larger-scale test and examine whether the development of writing influences properties of syntax. Such an effect could manifest in two distinct domains: (1) variations between speech and writing, where individuals may employ differing levels of syntactic complexity (a synchronic effect), and (2) the evolution of syntax over time, reflecting changes in grammatical structure when written styles influence norms of language even in speech (a diachronic effect). To explore these effects, we utilize two types of data: corpus data from the Universal Dependencies treebanks [34] and a dataset of annotated syntactic constructions designed to assess the influence of writing on syntactic evolution (Fig. 1).

**Figure 1.**
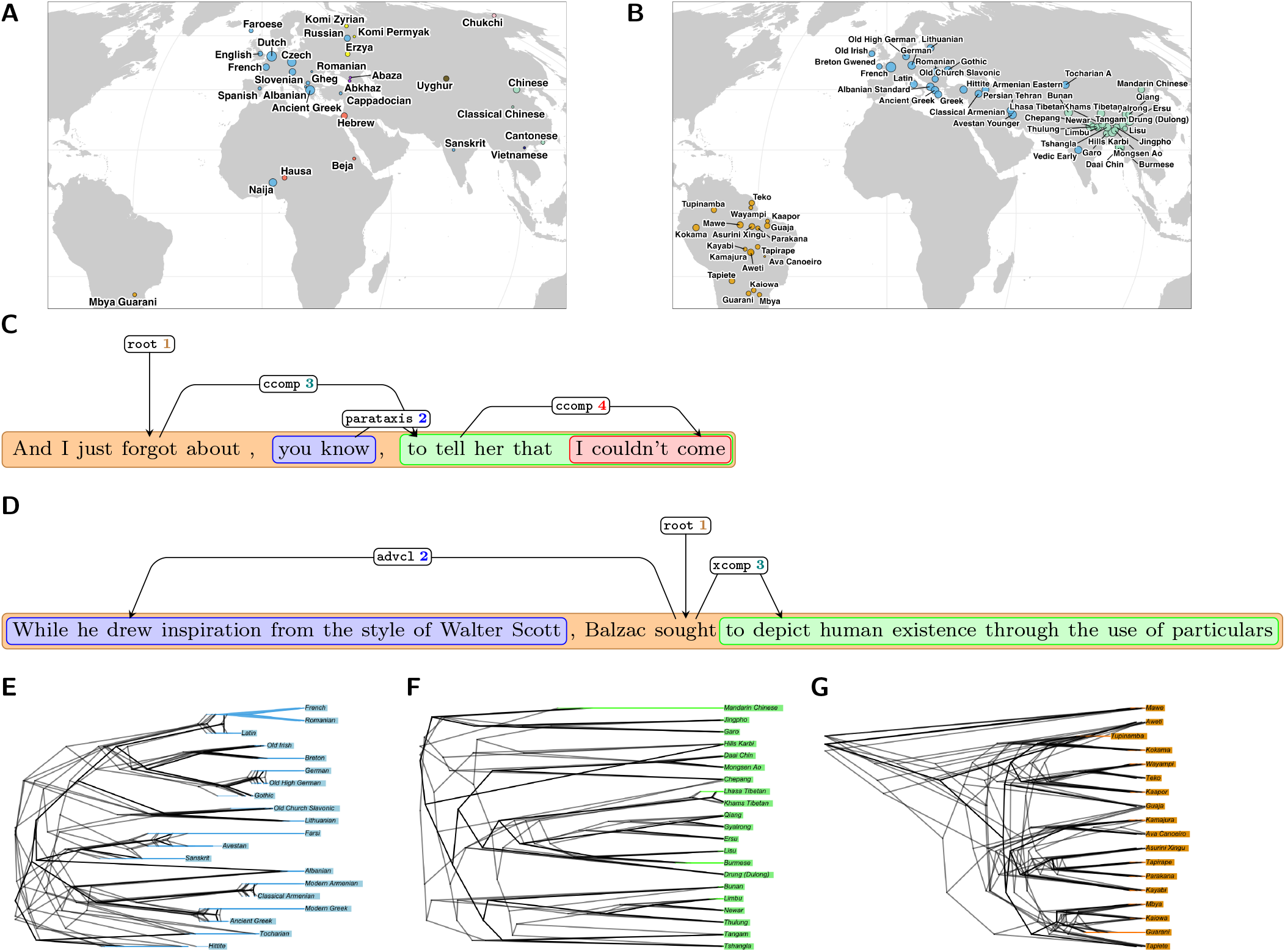
**A–B:** Geographic distribution of languages in the Universal Dependencies dataset (**A**) and in the syntactic construction dataset (**B**). Colors indicate language families, and point size corresponds to the number of sentences/constructions in each corpus. **C–D:** A spoken English sentence (**C**) and a sentence from the English Wikipedia (**D**) both with annotation of the syntactic relations between the clauses that constitute the sentence; dependent clauses are enclosed in colored boxes. **E–G:** Example of phylogenetic trees painted with two regimes. The black segments represent the “non-writing” regime, while the colored segments denote the “writing” regime. The length of the latter corresponds to the earliest attestation of a written document in each branch. The model was fitted on a sample of trees, with random noise added to the regime lengths to account for uncertainty regarding the onset of writing.

## Results

### Language Use

We used the Universal Dependencies corpus [35], which offers syntactic annotations for over 150 languages within a cross-linguistically consistent framework. Our analysis is based on 99,834 sentences drawn from three balanced genres: ‘Spoken’ (31,277 sentences), ‘Wikipedia’ (33,454), and ‘Fiction’ (35,103). These sentences represent 30 languages across 10 language families, including several non-LOL (non-”Literate, Official with Lots of users”) languages [36] such as Komi, Abaza, and Erzya. The geographic distribution of these languages is shown in Fig. 1A.

The ‘Spoken’ genre includes data from treebanks containing natural, spontaneous speech. These features are evident in the sentence shown in Fig. 1C, where hesitation and informal discourse structure illustrate the spontaneous nature of spoken language. In contrast, the ‘Wikipedia’ and ‘Fiction’ genres reflect more formal, edited language. For instance, the sentence shown in Fig. 1D (‘Wikipedia’ genre) exemplifies information-dense exposition with complex structure. While these examples align with the hypothesized impact of writing on syntax, the difference may be limited to English and may not replicate in other languages.

To assess syntactic complexity at scale, we used two metrics: the number of clauses per sentence and the maximum clausal path length. A clause is defined as any syntactic unit headed by a node with one of the following Universal Dependencies dependency relations: `root`, `acl`, `advcl`, `ccomp`, `csubj`, or `xcomp`. The maximum clausal path length captures the longest dependency path from the root to a terminal node, counting only these clausal nodes. Fig. 1C illustrates these metrics: the sentence contains four clauses — two embedded (“to tell her that”, “I couldn’t come”), one parenthetically inserted one (“you know”), and one main clause — while the maximum clausal path length is three, passing through `root`, `ccomp 3`, and `ccomp 4`. Clauses under `parataxis`, being concatenated rather than embedded, are excluded from the path length but included in clause counts; the overall distribution of these values is shown in Fig. S1.

We analyzed the data using a hierarchical Bayesian model that accounts for both phylogenetic and areal structure (see *Methods*). Model comparison indicated that a negative binomial likelihood provided the best fit for modeling the number of clauses (ΔELPD = 4621.4, SE = 174.9), while a Poisson likelihood was preferred for modeling clausal path length (ΔELPD = 2.9, SE = 0.6). Fig. 2B presents the average marginal effects [37] of genre on the number of clauses. All pairwise comparisons between genres center closely around zero, with already the 50% highest posterior density intervals (HPDI) overlapping the null effect line. This indicates that genre does not meaningfully predict the number of clauses per sentence. Similarly, Fig. 2A shows the marginal effects for clausal path length, revealing the same pattern: no significant differences across genres.

**Figure 2.**
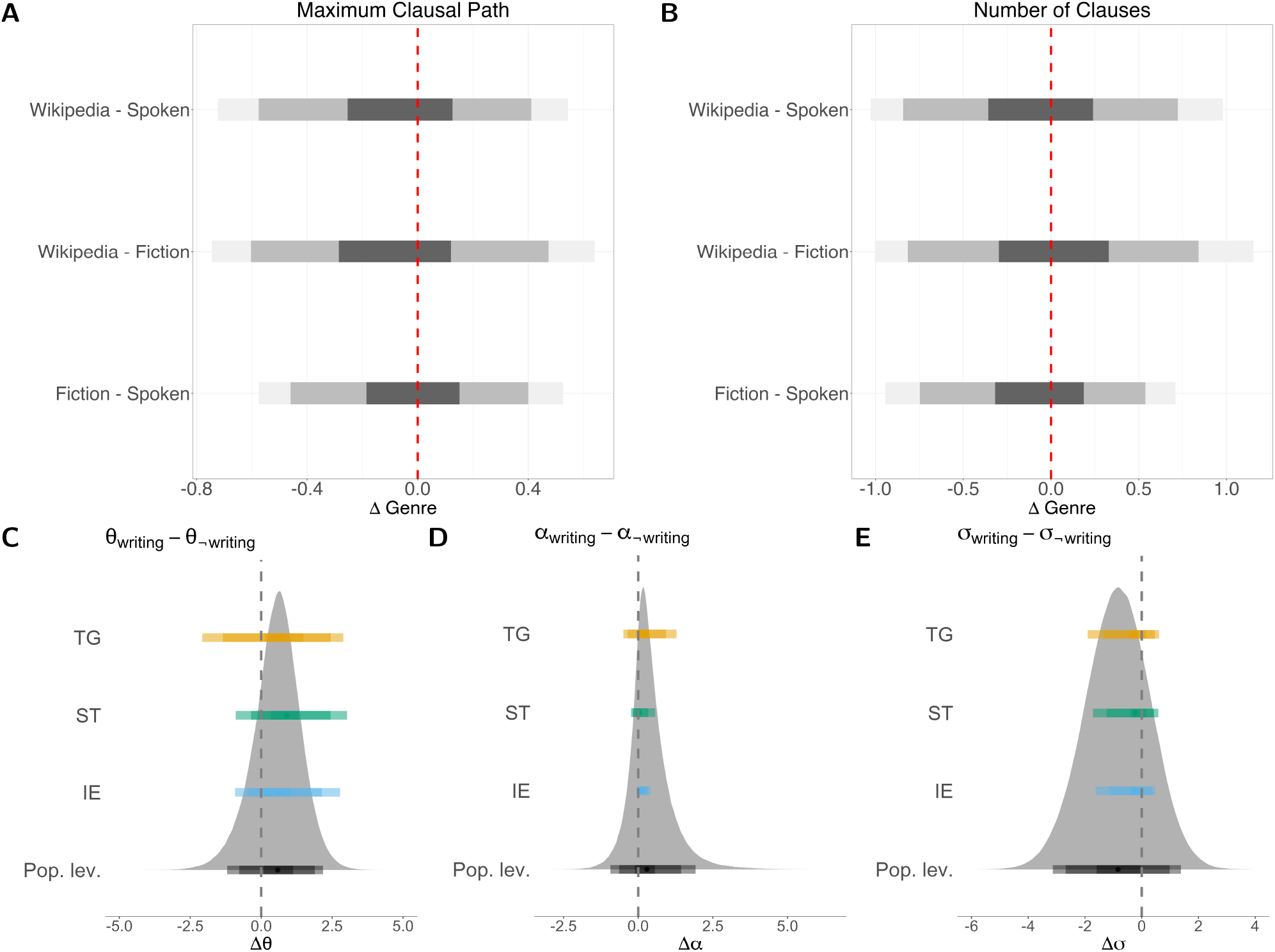
**A–B:** Marginal effects of genre on the maximum clausal path (**A**) and the number of clauses (**B**). **C–E:** Differences in parameter values in the evolution of syntactic hierarchy between the two regimes before vs after the introduction of writing. The density distribution in the background represents the differences in baseline (population-level) values, while the intervals in the foreground correspond to differences specific to each language family. All plots display highest posterior density intervals (HPDIs) at 50%, 89%, and 95% levels.

Random effects — accounting for phylogenetic similarity, areal proximity, and treebank-specific variation — likewise showed no substantial influence. Their 95% HPDI all included zero, and their standard deviations were consistently low (0.097–0.101), indicating minimal additional variance explained by these group-level factors (Fig. S6).

Taken together, the results suggest that syntactic complexity, as measured by clause count and maximum clausal path length, is not systematically influenced by genre in the Universal Dependencies data. Contrary to most case studies of individual languages, these metrics show no robust genre-related differences across the 30 languages analyzed (for detailed parameter values, cf. the Supplementary Information, Section S3.1).

### Language Grammar

While these findings suggest no direct impact of genre on individual production, the invention of writing might have biased the long-term evolution of grammar towards a more hierarchical structure.

To assess this possibility, we defined hierarchy in grammar as a partially ordered set and quantified it by counting the number of structural asymmetries that underlie the partial ordering in the combination of clauses. For instance, in the sentence “By getting the right people, we can do it”, three asymmetries emerge: (1) the presence of “by,” which presupposes a main clause; (2) the omission of the subject, indicating coreference with the second clause; and (3) the lack of tense and aspect markers, indicating inheritance from the second clause. By contrast, the sentence “We get the right people, we can do it” shows no structural asymmetry and, therefore, no evidence of hierarchy. As the example shows, the difference in form does not necessarily imply a difference in the message that is conveyed.

We coded nearly 700 clause-combining constructions for these asymmetries in 59 languages spanning three language families: Indo-European (IE), Sino-Tibetan (ST), and Tupí-Guaraní (TG) (see Fig. 1B for the complete list). These families were chosen for their diversity in writing traditions and population sizes, factors often linked to variation in clause combination [8]. For coding, we relied on a set of 18 asymmetry features reflecting diverse structural properties [21, 38] such as the presence of nominalization, the explicitness of linking elements, and syntactic or semantic overlap between clauses (Fig. S2-S4 show the distribution of the asymmetry features’ values across the languages in the data).

We modeled the evolution of logit-transformed binary asymmetries using an Ornstein-Uhlenbeck (OU) process [39, 40, 41] with two regimes: “non-writing” and “writing” (Fig. 1E–1G). The OU process captures the dynamics of continuous traits under stabilizing selection. It is defined by three parameters: the stationary trait value (*θ*), the strength of selection of this value (*α*), and the rate of stochastic fluctuations around it (*σ*). In other words, *θ* represents the long-term value toward which the trait evolves, *α* reflects the intensity of attraction to *θ*, and *σ* measures random variation.

We captured the OU process in a hierarchical Bayesian model and marginalized over phylogenetic uncertainty by fitting the model to a sample of approximately 50 trees per language family. The model included population-level parameters for the OU process (*θ*_*µ*_, *α*_*µ*_ and *σ*_*µ*_) as well as grouping-level values for each language family (*θ*_{IE,ST,TG}_, *α*_{IE,ST,TG}_, *σ*_{IE,ST,TG}_), allowing us to estimate long-term trait values at different phylogenetic scales. The length of the “writing” regime was scaled to the earliest attestation of writing in each branch, with added stochastic noise to account for uncertainty in the onset of writing.

While the posterior distributions of the OU parameters suggest slightly more asymmetries in written than non-written regimes, the evidence is weak. All differences between the parameter estimates in the “writing” minus the “non-writing” regime include 0 in their 95% HPDI (Δ*θ*_*µ*_ = [− 1.19, 2.19]), and the probability of a positive difference (Δ*θ*_*µ*_ *>* 0) is only 77% (Fig. 2C–2E). Thus, the two regimes appear to exhibit similar optimal levels of hierarchy.

Similarly, while we observed a small trend toward higher selection strength (*α*) and lower stochasticity (*σ*) in writing regimes, the evidence remains weak. The 95% HPDIs for both differences include zero (Δ*α*_*µ*_ = [−0.932, 1.92]; Δ*σ*_*µ*_ = [−3.14, 1.37]), and the probability of a positive or negative difference remains below 90% (Δ*α*_*µ*_ *>* 0 in 74% of samples, and Δ*σ*_*µ*_ *<* 0 in 77% of samples). This trend is only more pronounced in the Indo-European family: while the 95% HPDI for Δ*α*_IE_ still includes zero ([−0.06, 0.4]), the probability that Δ*α*_IE_ *>* 0 increases to 95%; additional details and visualizations are provided in Section S3.2.

## Discussion

Neither analysis revealed evidence that writing influences the amount of hierarchy or syntax complexity. For language use, the average marginal effects of written genres (“Fiction” and “Wikipedia”) did not differ decisively from those of spoken language. Similarly, for language grammar, the *θ* parameters of the two regimes showed no notable differences, with the posterior distribution of differences centered around zero. Contrary to the expectations of individual case studies, our findings therefore suggest that writing does not affect the degree of syntactic complexity in a language.

Our results suggest that the findings of a positive effect noted earlier are likely specific to individual languages and do not reflect a common or universal mechanism. In other words, the amount of hierarchy in language appears to vary in response to cultural ideas about ‘proper’ or ‘good’ speech, which permeate literate and preliterate societies worldwide [42, 43]. The contrasts that have been previously noted between written and spoken expression, whether within or between languages, might relate more to a specific cultural idea of expressive style than to direct effects of modality.

However, there are tentative indications of a writing-related small increase of *α* and decrease of *σ* parameters in Indo-European(Pr(Δ*α*_IE_ *>* 0) = 0.95, Pr(Δ*σ*_IE_ *<* 0) = 0.80). These differences may suggest a normativizing influence of writing. Since the effect is limited to one language family, the likely reason is again not writing per se but culturally specific ideas that yield a more regulated and uniform style of hierarchical expression.

While we do not demonstrate the absence of writing-related effects, our findings reflect an absence of evidence for such effects under the present analytical conditions. Limitations remain, notably the reliance on corpus data, which may not fully capture informal language use, and a sample that, while substantially larger than in previous studies, remains restricted in scope. Future research should aim to expand coverage across language families and communication modalities, and explore more deeply the sociocognitive mechanisms through which writing may influence grammatical change.

## Methods

All data and scripts used in this study can be found in the [Zenodo repository](https://doi.org/10.5281/zenodo.16107637).

### Universal Dependencies Data

The metadata were obtained by scraping the corpora available on the [Universal Dependencies website](https://universaldependencies.org/). Only corpora with clearly identified genres were included in the analysis. To compare spoken and written data with a comparable number of sentences, we selected the two written genres with the closest number of sentences to the spoken data. This resulted in three genres: Spoken (31,277 sentences), Wikipedia (33,454 sentences), and Fiction (35,103 sentences). The languages (with glottocodes [44]) and corresponding treebanks included in each genre are listed below:

- Spoken:

- Abaza (abaz1241): Abaza-ATB
- Beja (beja1238): Beja-NSC
- Cantonese (cant1236): Cantonese-HK
- Chinese (mand1415): Chinese-HK
- Chukchi (chuk1273): Chukchi-HSE
- English (stan1293): English-ESLSpok
- French (stan1290): French-ParisStories, French-Rhapsodie
- Albanian Gheg (gheg1238): Gheg-GPS
- Hausa (haus1257):Hausa-NorthernAutogramm, Hausa-SouthernAutogramm
- Komi Zyrian (komi1268): Komi_Zyrian-IKDP
- Naija (nige1257): Naija-NSC
- Slovenian (slov1268): Slovenian-SST
- Spanish (stan1288): Spanish-COSER
- Vietnamese (viet1252): Vietnamese-TueCL
- Wikipedia:

- Albanian Tosk (alba1268): Albanian-TSA
- Chinese (mand1415): Chinese-GSDSimp
- Dutch (dutc1256): Dutch-LassySmall
- Faroese (faro1244): Faroese-OFT
- Hebrew (hebr1245): Hebrew-IAHLTwiki
- Russian (russ1263): Russian-GSD
- Fiction:

- Abkhaz (abkh1244): Abkhaz-AbNC
- Ancient Greek (anci1242): Ancient_Greek-Perseus
- Cappadocian (capp1239): Cappadocian-TueCL
- Classical Chinese (lite1248): Classical_Chinese-TueCL
- Czech (czec1258): Czech-FicTree
- Erzya (erzy1239): Erzya-JR
- Komi Permyak (komi1269): Komi_Permyak-UH
- Komi Zyrian (komi1268): Komi_Zyrian-Lattice
- Mbya Guarani (mbya1239): Mbya_Guarani-Dooley
- Romanian (roma1327): Romanian-ArT
- Sanskrit (sans1269): Sanskrit-UFAL
- Uyghur (uigh1240): Uyghur-UDT

To compute the two measures of subordination — total clause count and maximum clausal path — we focused on dependency relations (denoted by `deprel`) that specifically marked clauses. These relations included:

- `root` (the root of the sentence)
- `acl` (adnominal clause)
- `advcl` (adverbial clause)
- `ccomp` (clausal complement without an obligatorily controlled subject)
- `csubj` (clausal syntactic subject)
- `parataxis` (clauses placed side-by-side without explicit coordination or subordination)
- `xcomp` (clausal complement with an obligatorily controlled subject)

The relation `parataxis` was only included in the calculation of the total clause count, as paratactic clauses are not, by definition, embedded within one another.

The total clause count was computed by summing all occurrences of these relations within a sentence. To calculate the maximum clausal path, all dependency paths from the root of a sentence to its terminal nodes were traced, and the longest path containing the specified dependency relations was identified. By definition, the total clause count always exceeded the maximum clausal path.

Additional phylogenetic and geographic information used to enrich the dataset was obtained from Glottolog [44] and AUTOTYP [45].

### Clause Combining Data

The dataset comprised a total of 59 languages from three language families (glottocodes in parentheses)

- Indo-European (IE): Modern Standard German (stan1295), Old High German (oldh1241), Gothic (goth1244), Old Irish (oldi1246), Breton (bret1244), Modern French (stan1290), Romanian (roma1327), Latin (lati1261), Old Church Slavonic (chur1257), Lithuanian (lith1251), Modern Armenian (nucl1235), Classical Armenian (clas1256), Modern Greek (mode1248), Ancient Greek (anci1242), Albanian (alba1268), Modern Persian (fars1255), Avestan (aves1237), Vedic (vedi1234), Tocharian (tokh1241), Hittite (hitt1242).
- Sino-Tibetan (ST): Qiang (qian1264), Gyalrong (jiar1240), Ersu (ersu1241), Lisu (lisu1250), Burmese (nucl1310), Lhasa (utsa1239), Khams (kham1282), Drung (dulo1243), Limbu (limb1266), Newar (newa1247), Thulung (thul1246), Bunan (gahr1239), Tangam (tang1377), Tshangla (tsha1245), Chepang (chep1245), Daai Chin (daai1236), Mongsen (mong1332), Hills Karbi (karb1241), Jingpho (jing1260), Garo (garo1247), Mandarin (mand1415).
- Tupí-Guaraní (TG): Tapirape (tapi1254), Xingu (xing1248), Parakana (para1312), Kayabi (kaya1329), Kamajura (kama1373), Canoe (avac1239), Mbya (mbya1239), Guarani (jopa1240), Kaiowa (kaiw1246), Tapiete (tapi1253), Wayampi (waya1270), Teko (emer1243), Kaapor (urub1250), Tupinamba (nhen1239), Kokama (coca1259), Guaja (guaj1256), Aweti (awet1244), Mawe (sate1243).

The dataset comprised a total of 763 constructions: 317 in IE, 335 in ST, and 111 in TG. On average, there were 16 constructions per language in IE and ST, and 6.17 constructions per language in TG. The standard deviation of constructions per language was 5.1 for IE, 5.73 for ST, and 3.28 for TG.

A syntactic structure was classified as a distinct construction if it exhibited at least one type of asymmetry distinguishing it from similar constructions.

The sampled constructions were analyzed according to 18 asymmetry features, each capturing specific syntactic relationships between the clauses within the constructions [38]:

- IDs 1-6: Agreement and tense-aspect-mood properties of the clauses.
- IDs 7-12: Degree of nominalization in the construction.
- ID 13: Nature of syndesis between clauses.
- ID 14: Relativization.
- IDs 15-18: General properties of clause independence, such as negation scope and clause order.

### Phylogenetic Regression

We modeled the number of clauses (`cl_number`) and maximum clausal path (`chain_len`) as count responses regressed on text genre, with random intercepts for `treebank`, geographical `area`, and phylogenetic relationships (`phylo`). Both models were implemented in a Bayesian framework using the `brms` [46] interface to Stan [47] in R [48].

We assumed Poisson and negative binomial likelihoods, depending on the dispersion properties of each response variable, and tested alternative model specifications using leave-one-out cross-validation (LOO-CV). For both outcomes, model comparison included alternatives with and without spatial Gaussian Processes (GPs) and with area random effects derived from AUTOTYP [45]. For clause counts, the model with areas as random effects was preferred; for maximum clausal path, model fits were comparable. To maintain consistency, we used area random effects in both final models.

Random effects for phylogenetic structure were informed by a variance-covariance matrix based on shared nodes in Glottolog 5.1 [44], constructed via the `vcv.phylo()` function from the `ape` package [49].

We applied weakly informative priors: *N* (0, 2) for intercepts and fixed effects, and Half-Normal(1, 0.1) for standard deviations of group-level effects. Models were estimated via MCMC with 4 chains of 8000 iterations each (4000 warm-up), with `adapt_delta = 0.9` to reduce divergent transitions.

Marginal effects were extracted using the `marginaleffects` package [37]. Full model formulas and diagnostics are presented in the SI, sections S2.1 and S3.1. Likewise, posterior predictive checks from the models are found in the SI, section S4.

### Ornstein-Uhlenbeck Model

To model the evolution of syntactic asymmetry features across Indo-European, Sino-Tibetan, and Tupí-Guaraní languages, we implemented an Ornstein-Uhlenbeck process following [50], with regime-specific variation along phylogenetic branches. Regimes were defined by the presence or absence of writing, and allowed intercepts (*θ*), evolutionary rates (*α*), and variances (*σ*) to vary across language families and features.

The model incorporated matrices *W* and *V* to encode expected states and phylogenetic covariances, respectively, based on branch-specific regime information and shared ancestry. Language-specific deviations from expected values were modeled as multivariate normal with covariance matrix *V*, allowing the process to flexibly accommodate heterogeneity across tips.

Hierarchical priors allowed partial pooling across families and features. All hyperparameters — means and scales for *θ, α*, and *σ* — were assigned weakly informative *N* (0, 1) priors while feature-level variability (*σ*_feature_) received a broader *N* (0, 1.5) prior. Latent variables and residuals were modeled as standard normal. The model was estimated in Stan [51] using 4 chains of 10,000 iterations each (5,000 warmup), with `adapt_delta` set to 0.99 to reduce divergences. Full implementation details, including regime encoding and tree structure, are available in the SI, section S2.2.

## Acknowledgements

We are grateful to Mathias Jenny (Sino-Tibetan), Rik van Gijn (Tupí-Guaraní), and Paul Widmer (Indo-European) for their help with data collection and to Lothar Sebastian Krapp for advice on the mathematical definitions of hierarchy.

## Contributions

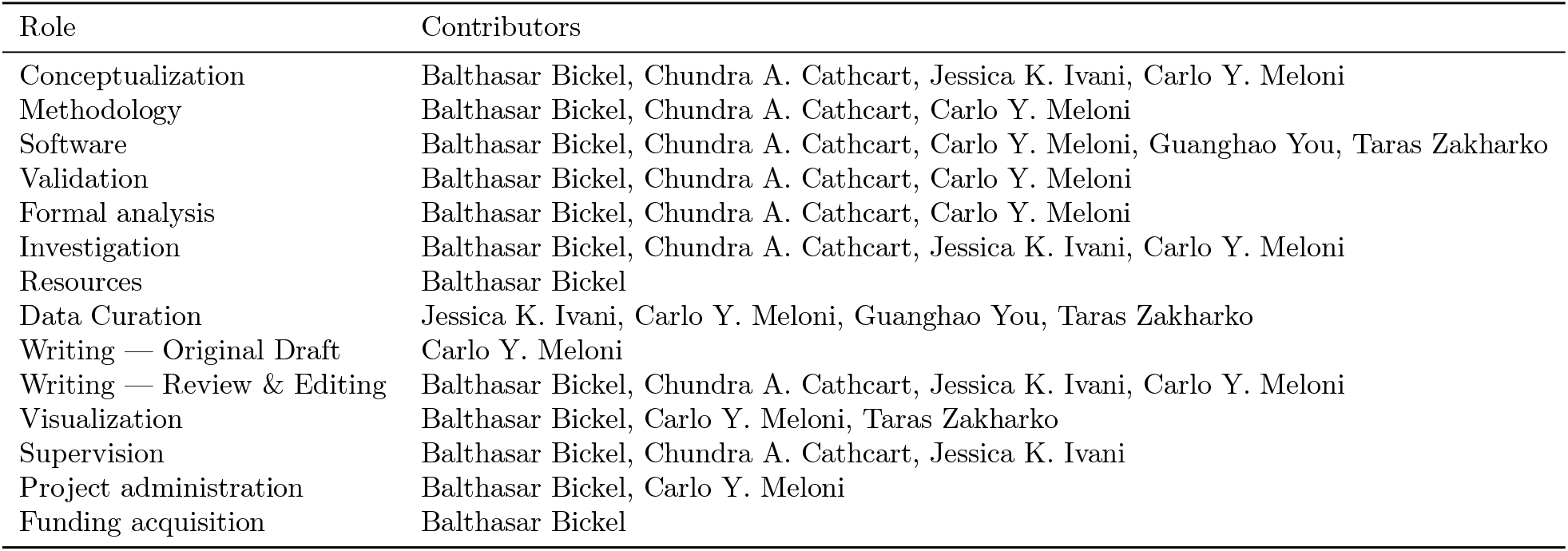

## Supplementary Information

**Figure S1.**
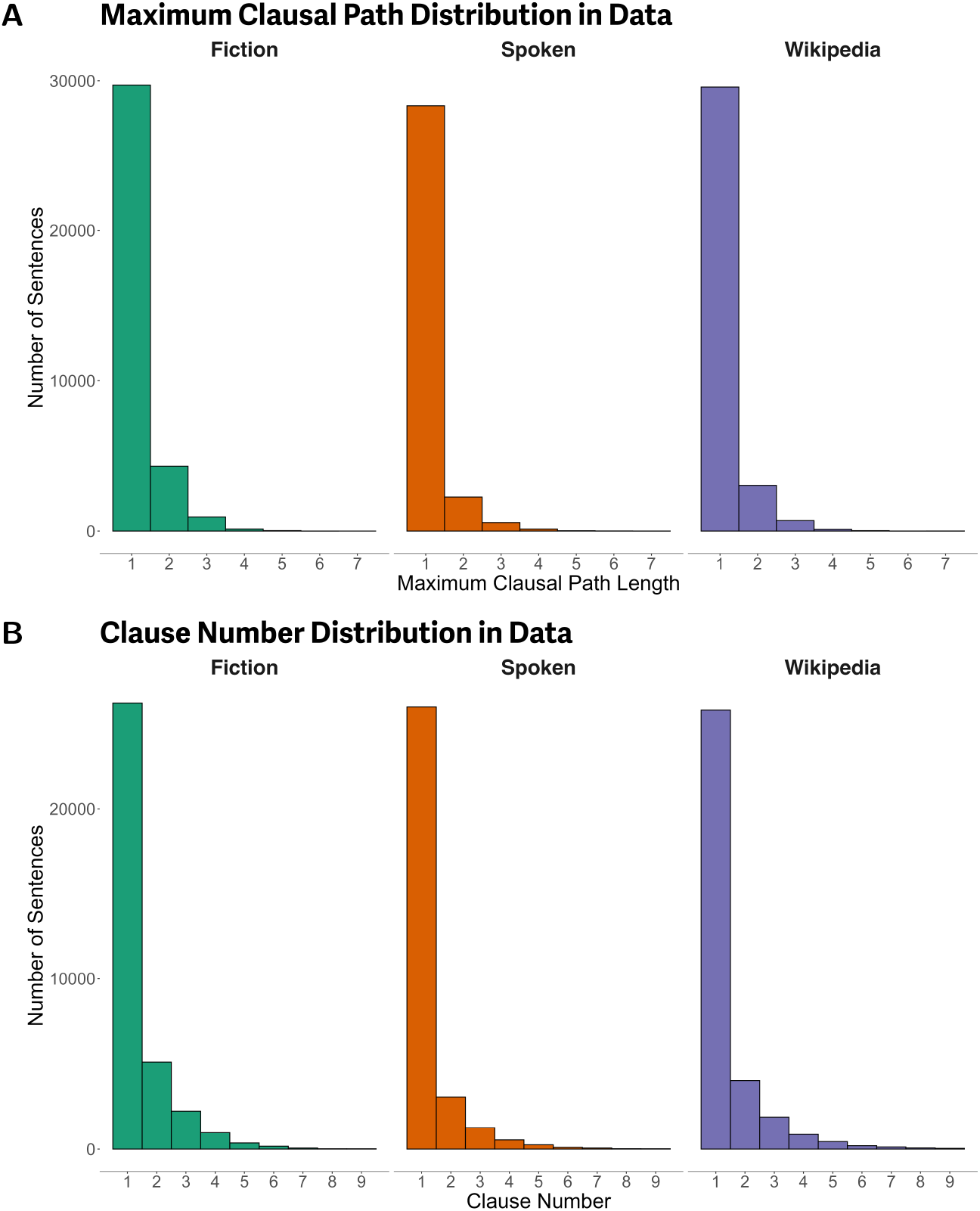
Distribution of sentence-level structural complexity in the language usage data. **A:** Distribution of the maximum clausal path length of sentences by genre. **B:** Distribution of the number of clauses per sentence by genre. **Note:** In Figure **B**, the x-axis has been truncated after the value 9 for clarity. The actual distribution has a longer tail with a small number of sentences containing high clause counts.

**Figure S2.**
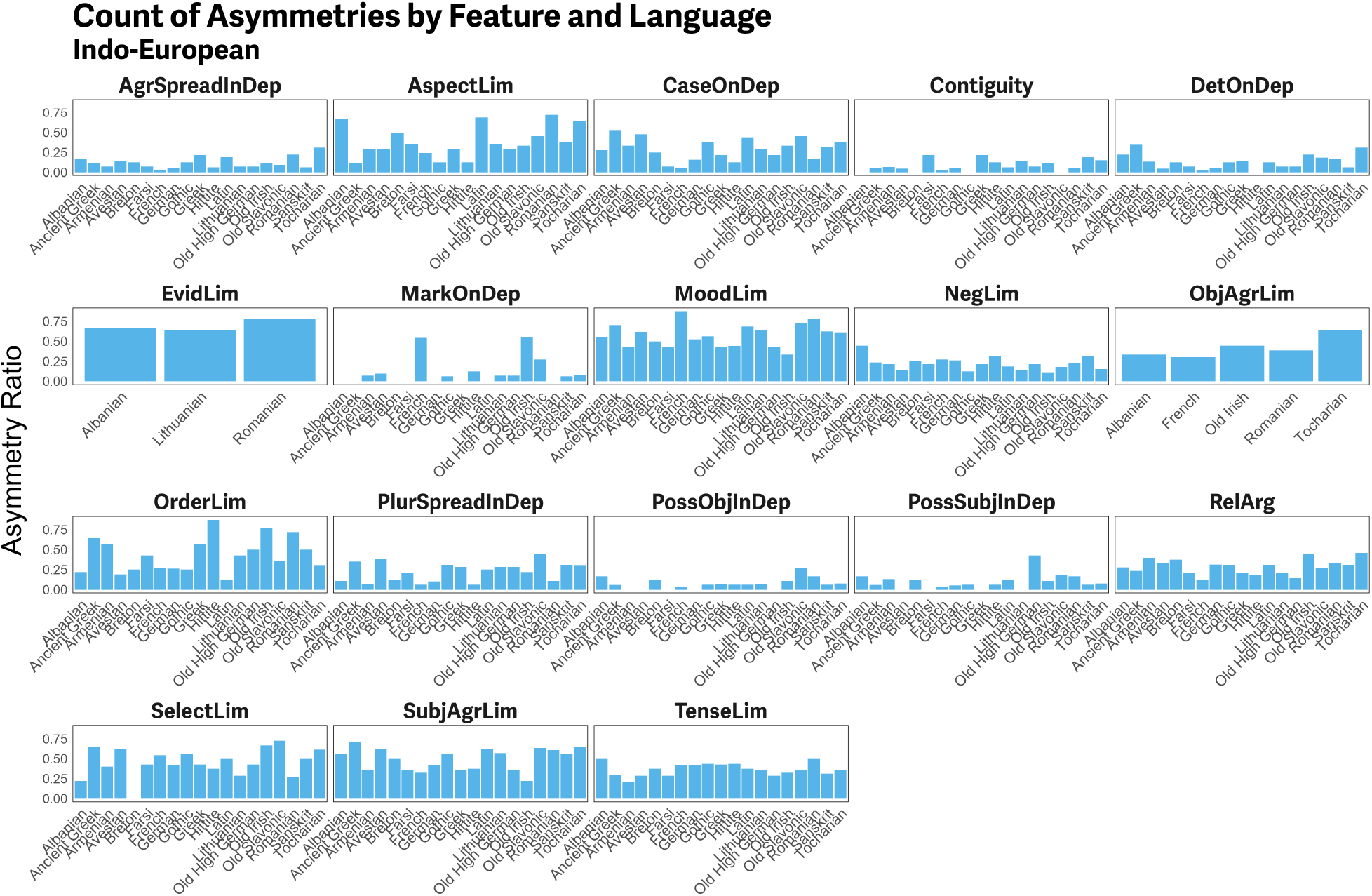
Distribution of data used in the language grammar analysis in Indo-European. The facets represent different asymmetry features. The y-axis shows the proportion of constructions in each language for which the asymmetry feature is present (i.e., equals 1), calculated as 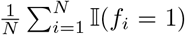, where *N* is the number of constructions in the language, *f*_*i*_ is the value of the feature for the *i*-th construction and II (·) is the indicator function (1 if true, 0 otherwise). The x-axis displays individual languages within each language family.

**Figure S3.**
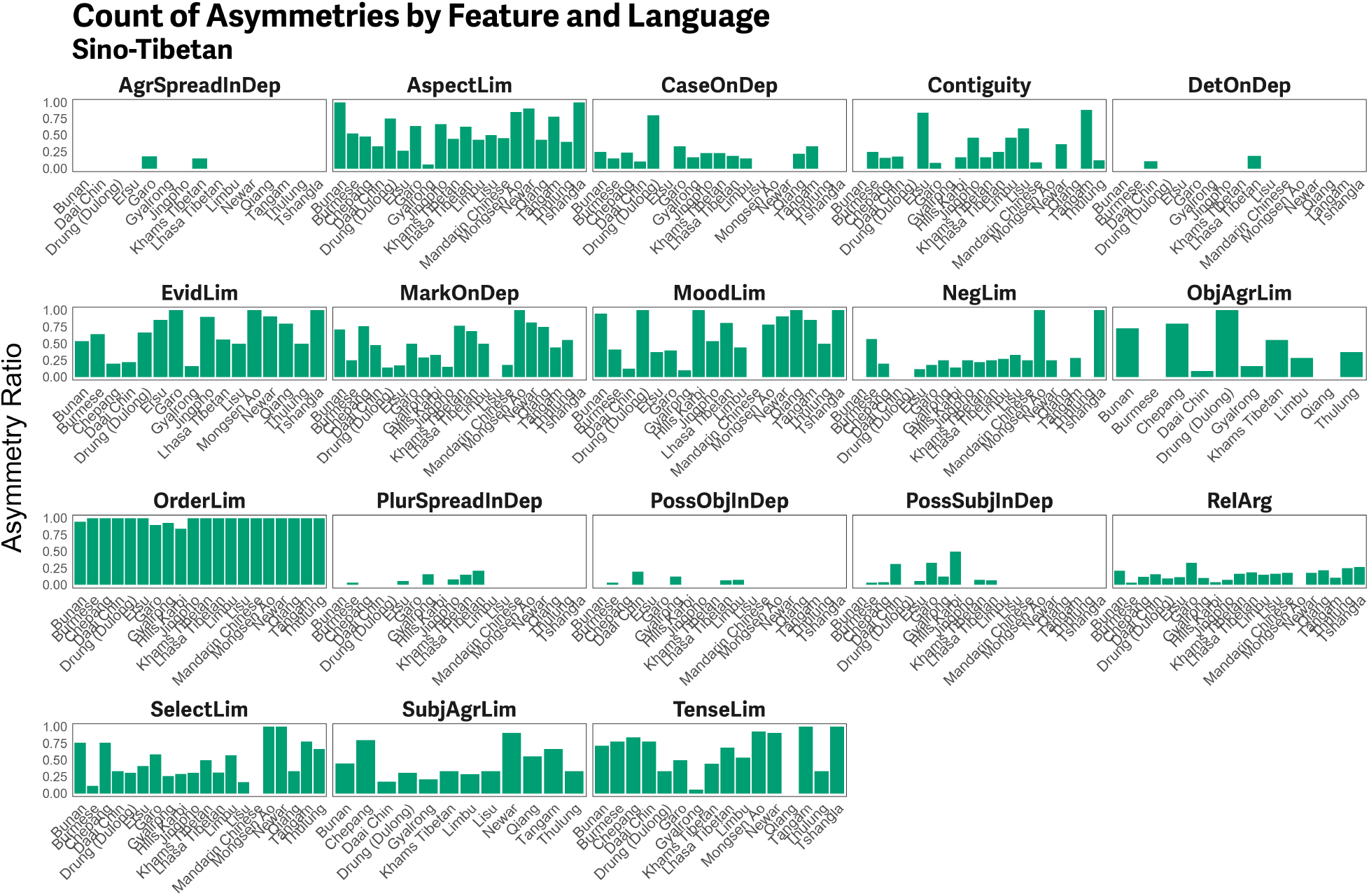
Distribution of data used in the language grammar analysis in Sino-Tibetan. Same conventions as in Figure S1.

**Figure S4.**
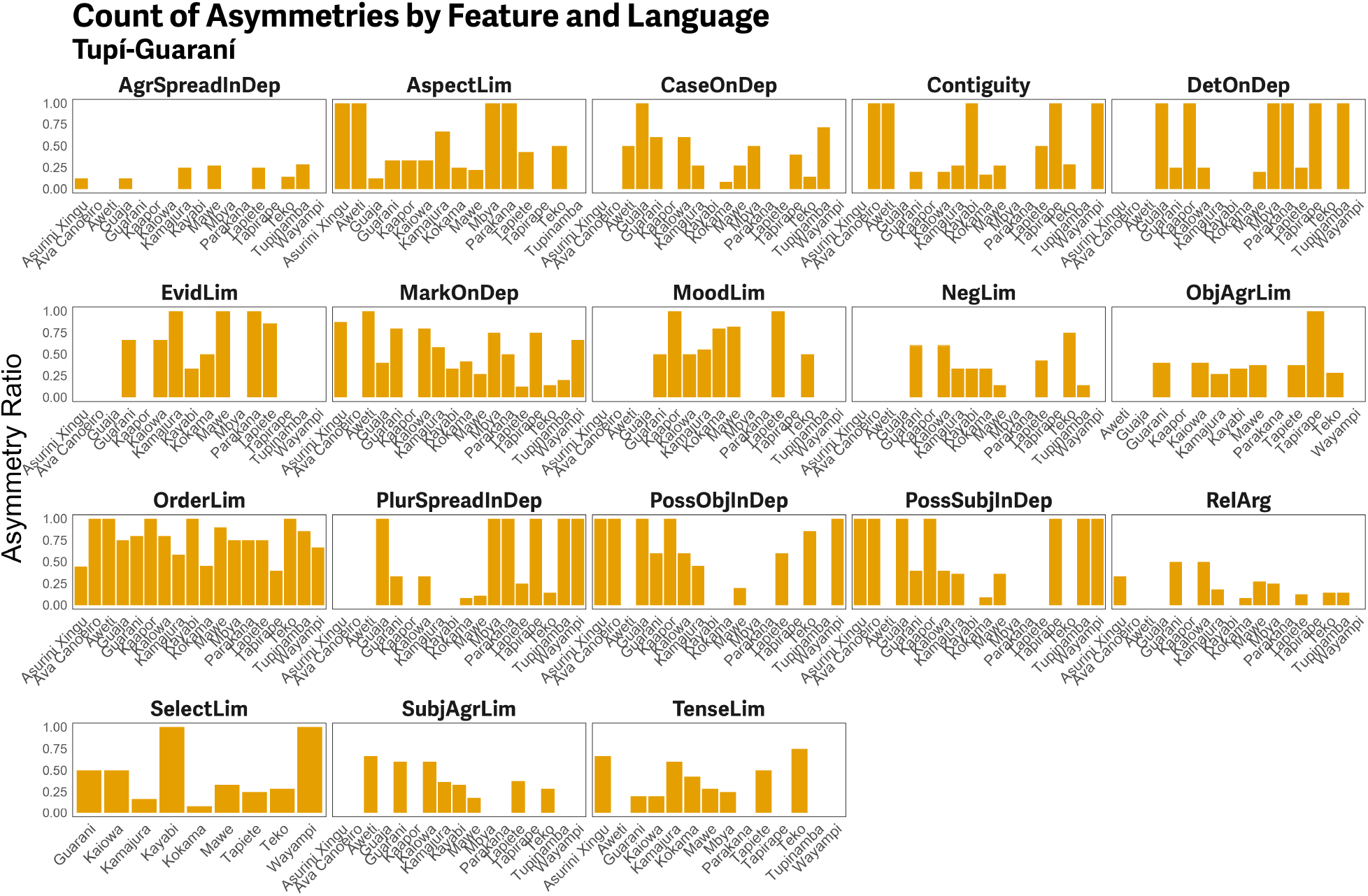
Distribution of data used in the language grammar analysis in Tupí-Guaraní. Same conventions as in Figure S1.

## Model Description

### Phylogenetic Regression

Two distinct models were implemented to analyze linguistic usage patterns, differing in their choice of response variable. The first model used the maximum clausal path as the response variable, while the second model examined the number of clauses. In both models, the response variable was modeled as a function of text genre, while accounting for additional variability arising from linguistic family relationships, geographical distribution, and treebank-specific idiosyncrasies. These models were implemented in Bayesian framework using the `brms` [46] interface to Stan [47] in R [48].

Let:

- *y*_*i*_ be the response variable for observation *i*
- *η*_*i*_ be the linear predictor
- *β*_0_ be the intercept
- *β*_*j*_ be the fixed effect for genre level *j*
- *a*_*a*[*i*]_ be the random intercept for geographical area
- *t*_*t*[*i*]_ be the random intercept for treebank
- *p*_*p*[*i*]_ be the phylogenetic random effect
- Σ_phylo_ be the phylogenetic covariance matrix based on shared ancestry
- *θ* be the shape (dispersion) parameter in the negative binomial distribution We define indicator variables for the genre levels:

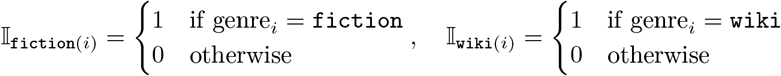

### Poisson Model for Maximum Clausal Path

The first model used maximum clausal path as the dependent variable and assumed it followed a Poisson distribution with a log link, appropriate for count data [52]. The fixed effect in this model was `genre`, a three levels: `spoken`, `fiction`, and `wiki`. The variable was treated as an ordered factor (`spoken` *<* `fiction` *<* `wiki`) to capture a hypothesized gradient of syntactic complexity across these genres. The random effects structure accounted for variability at three levels: `treebank`, geographical `area`, and phylogenetic relationships (`phylo`).

#### Likelihood

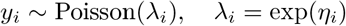

#### Linear Predictor

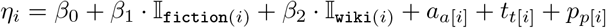

#### Random Effects

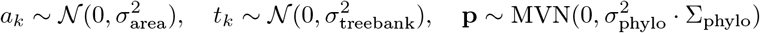

#### Priors

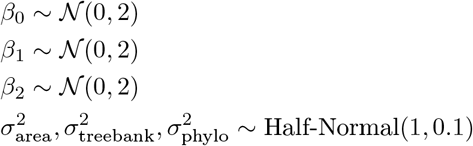

### Negative Binomial Model for Clause Number

The second model analyzed the count of clauses in a sentence, assuming a negative binomial distribution with a log link for the response variable. The fixed and random effects specifications mirrored those in the maximum catena length model. Specifically, `genre` was included as the primary fixed effect, and random intercepts for `treebank`, `area`, and phylogenetic relationships (`phylo`) were incorporated to capture structured variability.

#### Likelihood

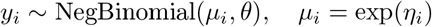

- *θ*: shape (overdispersion) parameter, such that variance = *µ*_*i*_ +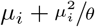

##### Linear Predictor

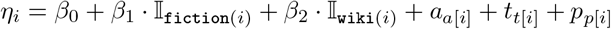

##### Random Effects

Same structure as in the previous model:

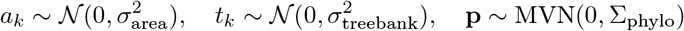

##### Priors

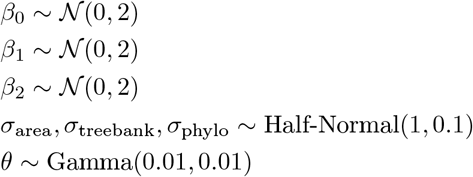

### Phylogenetic Covariance Matrix

Phylogenetic relationships were based on a variance-covariance matrix derived from Glottolog 5.1 [44], constructed by calculating the number of shared nodes between languages. This matrix was generated using the `vcv.phylo()` function from the `ape` package [49]:

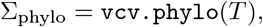

where *T* is the tree derived from Glottolog (with unit length). The resulting covariance structure aligned the model’s intercept correlations with phylogenetic proximity, ensuring that closely related languages exhibited more similar effects, while unrelated families were modeled as independent (with zero covariance). Languages were grouped at the microfamily level (the smallest genealogical groupings above the leaf nodes [53]) to provide a detailed and informative representation of their phylogenetic relationships.

### Model Comparison

We controlled for areal effects through two approaches: (i) we incorporated the area variable from AUTOTYP[45]as a group-level effect, and (ii) included a Gaussian Process (GP) term in the model using the longitude and latitude point estimates from Glottolog of each language. GPs are non-parametric methods designed to capture non-linear dependencies, where proximity between observations determines the strength of their mutual influence [54]. To account for areal effects in two dimensions, we used a two-dimensional GP to model spatial dependencies across the geographical distribution of languages. Assuming minimal contact between areas (taken from AUTOTYP), we added independent GPs for each macro-area to better reflect regional variation [53].

We conducted model comparison using leave-one-out cross-validation to determine which method best accounted for areal effects. For the clause number data, the model with areas as random effects was preferred, while for catena length, no significant differences were observed between the models. To maintain consistency and facilitate comparability, we retained random effects by area in both models.

### Computational Details

The models were estimated using Markov Chain Monte Carlo (MCMC) sampling, with four chains of 8000 iterations each, including 4000 warmup iterations. To improve convergence and minimize divergent transitions, the `adapt_delta` parameter was set to 0.9. The marginal effects were estimated based on the model results using the `marginaleffects` package [37].

### Ornstein-Uhlenbeck Model

The Ornstein-Uhlenbeck (OU) model implemented here is designed to capture the evolution of syntactic asymmetry features across three language families: Indo-European, Sino-Tibetan, and Tupí-Guaraní. The model accounts for phylogenetic relationships between languages and allows for varying evolutionary dynamics depending on the feature under consideration and the evolutionary regimes along the branches of the phylogenetic trees. This framework was inspired by the model outlined in [50].

For each syntactic feature *f* ∈ {1, …, *F*} and for each language *l* ∈ {1, …, *T*_*k*_} in family *k* ∈ IE, ST, TG, we model the proportion of syntactic constructions with the asymmetry feature as a binomial outcome. Each observation *i* ∈ {1, …, *N*_*k*_} corresponds to *n*_*i*_ trials with *y*_*i*_ successes. The success probability *p*_*i*_ is modeled on the logit scale as the sum of two components: a language-level predictor log *p*_lang,*l*[*i*]_ and a feature-specific intercept log *p*_feat,*f*[*i*]_. The likelihood for each observation is thus:

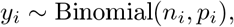

with

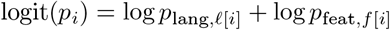

To account for evolutionary constraints and phylogenetic relationships, we model the language-level predictor log *p*_lang,*l*_ using an Ornstein–Uhlenbeck process. The evolutionary state of each language *l* is modeled as:

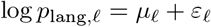

where *µ*_*l*_ is the expected evolutionary state based on ancestral trajectories, and *ε*_*l*_ is a residual deviation from this trajectory. The expectation *µ*_*l*_ is computed from a regime-weighted matrix *W*, such that:

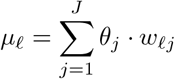

where *θ*_*j*_ is the evolutionary optimum associated with regime *j*, and *w*_*lj*_ encodes the contribution of regime *j* along the evolutionary path to language *l*, derived from start and end times along branches and their associated regimes. These weights are computed via the `gen_W` function [50], which integrates evolutionary decay and regime assignment along each branch using the OU kernel.

Residual deviations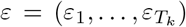 are modeled jointly as multivariate normal with mean zero and covariance matrix Σ, which is constructed to reflect shared evolutionary history:

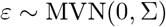

The matrix Σ is generated using the `gen_V` function [50], which incorporates phylogenetic distances and shared ancestry (via MRCA matrices) to define pairwise covariances under the OU process. The off-diagonal entries of Σ encode the expected covariance due to shared ancestry under different evolutionary rates *α*_*j*_ and variances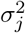per regime. The diagonal entries additionally include a small constant cons for numerical stability.

Each of the evolutionary parameters – optima *θ*_*j*_, rates *α*_*j*_, and variances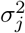 – is allowed to vary across families and is drawn from a shared set of hyperpriors, allowing for partial pooling. Specifically, for each family *k*, we define:

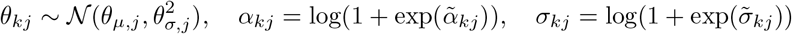

with latent variables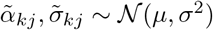, ensuring positivity of the rate and variance parameters through a softplus transformation.

Feature-specific intercepts *λ*_*f*_ = log *p*_feat,*f*_ capture baseline tendencies for integration and are modeled hierarchically:

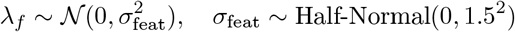

These intercepts are shared across families and added to the language-level predictors in the logit link function. The full predictive model thus integrates both family-specific evolutionary dynamics and feature-specific tendencies.

All parameters are given weakly informative priors to encourage regularization while allowing flexibility in posterior estimation. The latent z-parameters for hierarchical effects (e.g., *θ*_*z*_, *α*_*z*_, *σ*_*z*_) are all given standard normal priors. Residual terms *ε* are also drawn from a standard normal prior before being transformed via the Cholesky decomposition of Σ, ensuring a proper multivariate normal structure.

The model was implemented in Stan [51] and estimated using Markov chain Monte Carlo (MCMC) sampling with four chains, each running for 10,000 iterations (5,000 warmup). To improve convergence and reduce divergence issues in posterior sampling, the `adapt_delta` parameter was set to 0.99.

### Painting Phylogenetic Trees with Writing Regimes

To investigate the influence of writing systems on the evolution of languages, we introduced a two-regime model across phylogenetic trees representing different language families. These regimes distinguish between historical periods before and after the adoption of writing. Specifically, branches of the phylogenies were annotated with a “non-writing” regime from the root up to the point of writing adoption, and a “writing” regime thereafter.

The procedure began by sampling a subset of posterior trees for each family and pruning them to retain only the languages for which writing onset estimates were available. For each terminal branch (i.e., leading to a modern language), we estimated the point in time at which writing was adopted. These cut-points were derived from published historical and ethnographic sources specific to each language, and to account for uncertainty, small amounts of random variation were added using truncated normal distributions. This stochastic perturbation ensures that the timing of the regime shift is plausible but not treated as fixed.

Each selected tree was then modified by inserting a new internal node on the branch corresponding to a language’s writing onset. This involved splitting the original branch into two parts: the portion prior to the inserted node retained the “non-writing” regime, and the portion following the node was assigned to the “writing” regime. The location of the node along the branch was calculated in terms of time from the root, and adjusted to remain within the valid length of the original branch to preserve tree integrity.

After insertion, branches were “painted” according to their regime using discrete character states (e.g., “0” for non-writing and “1” for writing). This regime mapping enabled downstream comparative analyses to model trait evolution under different selective pressures. Trees with newly created singleton nodes (resulting from regime shifts near terminal tips) were further processed to standardize their structure, and all regime-mapped trees were saved for later use in OU model fitting.

## Parameter Posteriors

### Language Usage

#### Posterior Visualization

**Figure S5.**
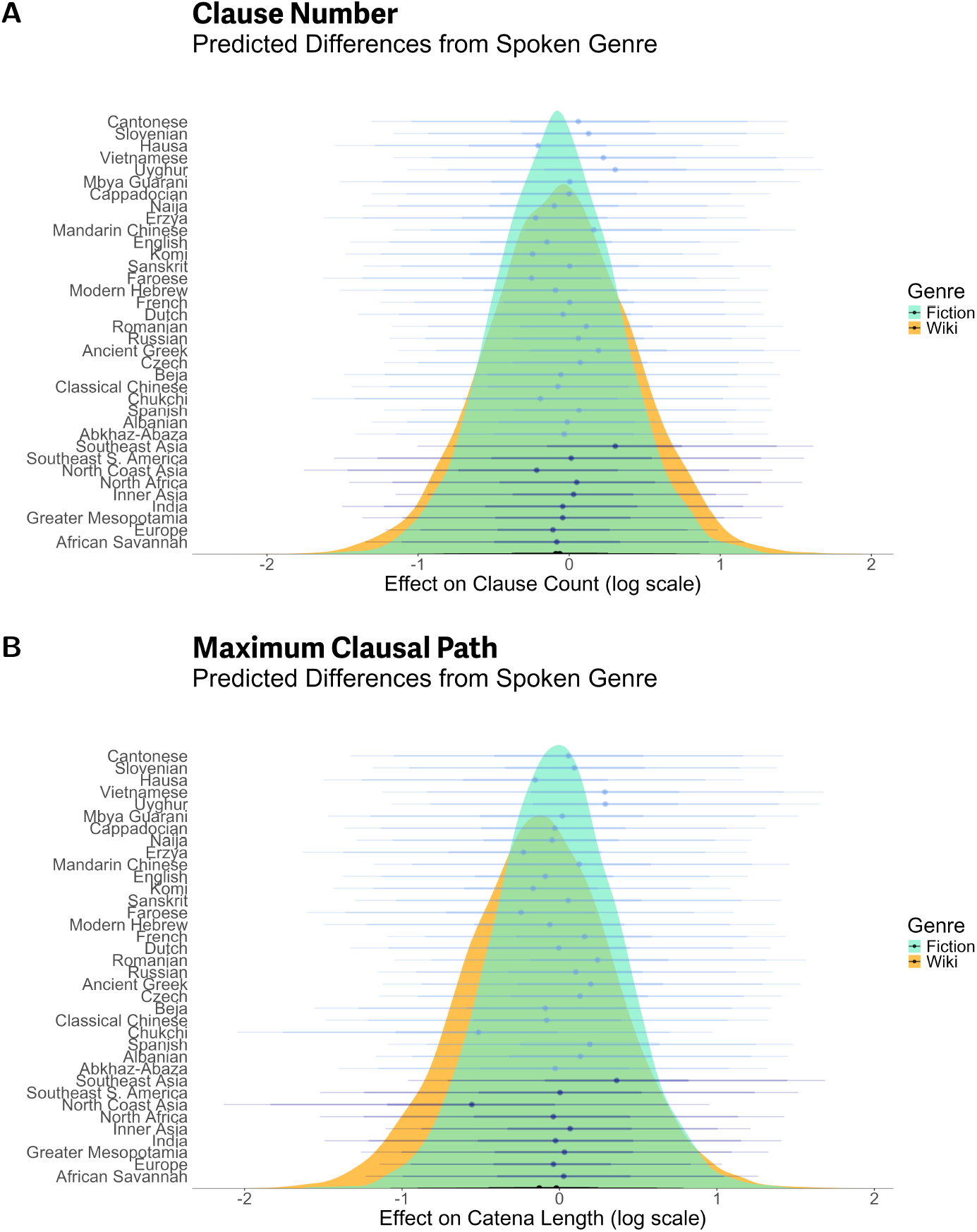
Posterior distributions of the random and fixed effects in the phylogenetic regression. The background distributions represent the posterior estimates for the fixed effect of *Genre*, expressed as predicted differences from the baseline category (‘Spoken’ genre). The foreground intervals indicate the random effects, with light blue representing phylogenetic effects and dark blue representing areal effects. **A:** Posteriors for the clause number model. **B:** Posteriors for the maximum clausal path model.

#### Parameter Values

**Table S1.** Multilevel hyperparameters for maximum clausal path model.

| Group | Parameter | Estimate | Est. Error | 2.5% CI | 97.5% CI | Rhat | Bulk ESS | Tail ESS |
| --- | --- | --- | --- | --- | --- | --- | --- | --- |
| **area** | sd(Intercept) | 0.96 | 0.10 | 0.76 | 1.16 | 1.00 | 29584 | 11190 |
| **phylo** | sd(Intercept) | 0.93 | 0.10 | 0.73 | 1.12 | 1.00 | 22312 | 11144 |
| **treebank** | sd(Intercept) | 0.83 | 0.10 | 0.64 | 1.03 | 1.00 | 14976 | 11105 |

**Table S2.** Regression coefficients for maximum clausal path model.

| Parameter | Estimate | Est. Error | 2.5% CI | 97.5% CI | Rhat | Bulk ESS | Tail ESS |
| --- | --- | --- | --- | --- | --- | --- | --- |
| Intercept | -0.77 | 0.51 | -1.76 | 0.23 | 1.00 | 9452 | 10956 |
| **genre** (fiction) | -0.02 | 0.40 | -0.80 | 0.77 | 1.00 | 7980 | 9862 |
| **genre** (wiki) | -0.13 | 0.48 | -1.07 | 0.79 | 1.00 | 7747 | 9758 |

**Table S3.**
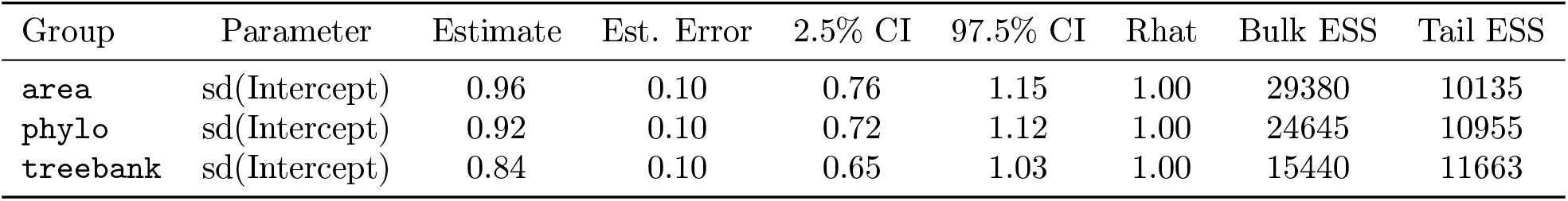
Multilevel hyperparameters for clause number model.

| Group | Parameter | Estimate | Est. Error | 2.5% CI | 97.5% CI | Rhat | Bulk ESS | Tail ESS |
| --- | --- | --- | --- | --- | --- | --- | --- | --- |
| **area** | sd(Intercept) | 0.96 | 0.10 | 0.76 | 1.15 | 1.00 | 29380 | 10135 |
| **phylo** | sd(Intercept) | 0.92 | 0.10 | 0.72 | 1.12 | 1.00 | 24645 | 10955 |
| **treebank** | sd(Intercept) | 0.84 | 0.10 | 0.65 | 1.03 | 1.00 | 15440 | 11663 |

**Table S4.** Regression coefficients for maximum clause number model.

| Parameter | Estimate | Est. Error | 2.5% CI | 97.5% CI | Rhat | Bulk ESS | Tail ESS |
| --- | --- | --- | --- | --- | --- | --- | --- |
| Intercept | -0.30 | 0.52 | -1.31 | 0.72 | 1.00 | 12070 | 11112 |
| **genre** (fiction) | -0.09 | 0.41 | -0.89 | 0.72 | 1.00 | 9892 | 10248 |
| **genre** (wiki) | -0.06 | 0.48 | -1.00 | 0.87 | 1.00 | 9719 | 10537 |
| shape | 1.78 | 0.03 | 1.72 | 1.83 | 1.00 | 33652 | 11350 |

#### Comparison of Parameter Uncertainty Across Model Components

**Figure S6.**
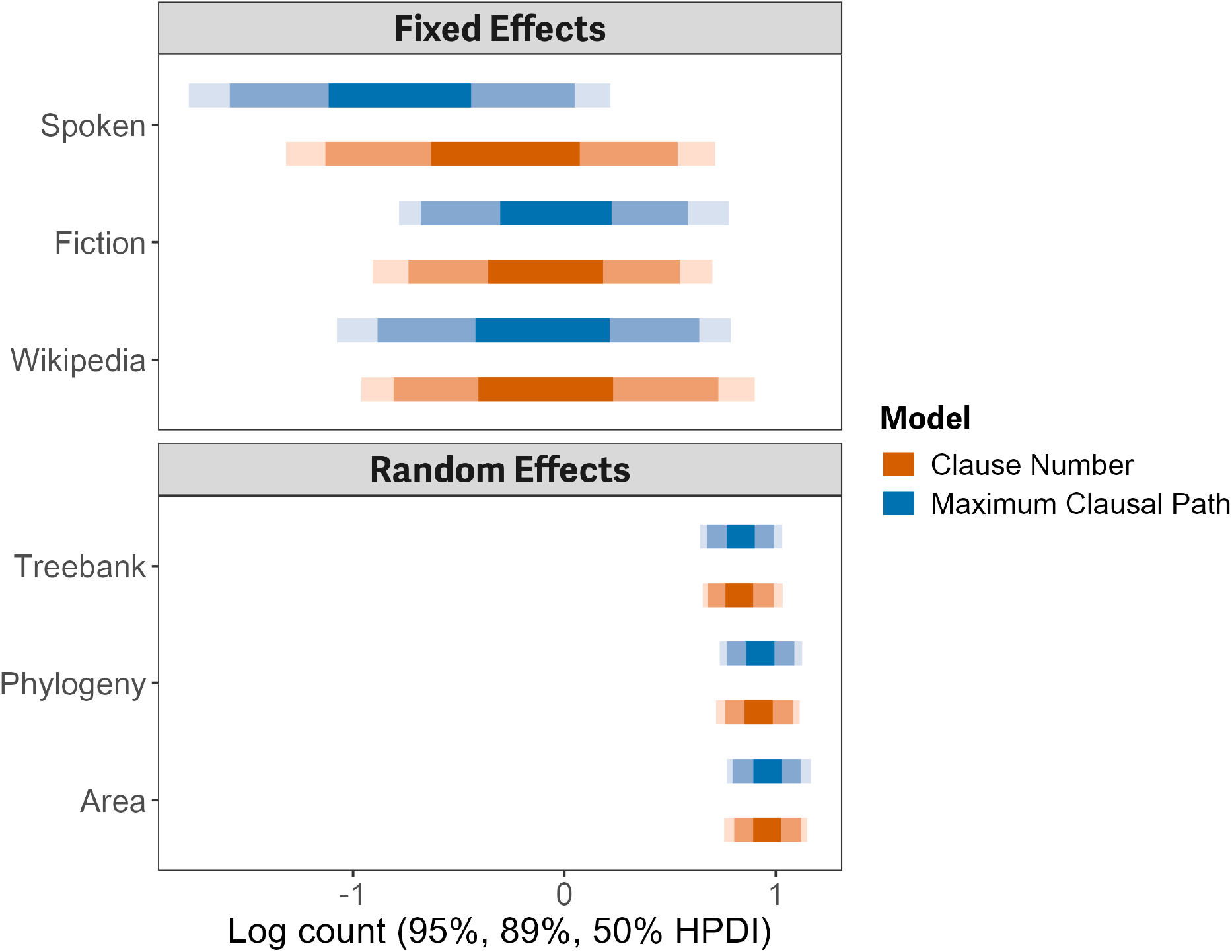
Comparison of the uncertainty in model parameters between fixed and random effects for two regression models: Clause Number and Maximum Clausal Path. The plot shows the posterior distributions of the fixed effects (“Spoken”, “Fiction”, “Wikipedia”) and standard deviations of the random effects grouped by area, phylogeny, and treebank. Across both models, fixed effects exhibit substantially greater uncertainty, as indicated by their wider HPDIs. In contrast, random effects display low standard deviations (range: 0.097– 0.101) and 95% HPDIs that consistently include zero, suggesting minimal additional variance explained by these group-level factors.

#### Language Grammar

**Figure S7.**
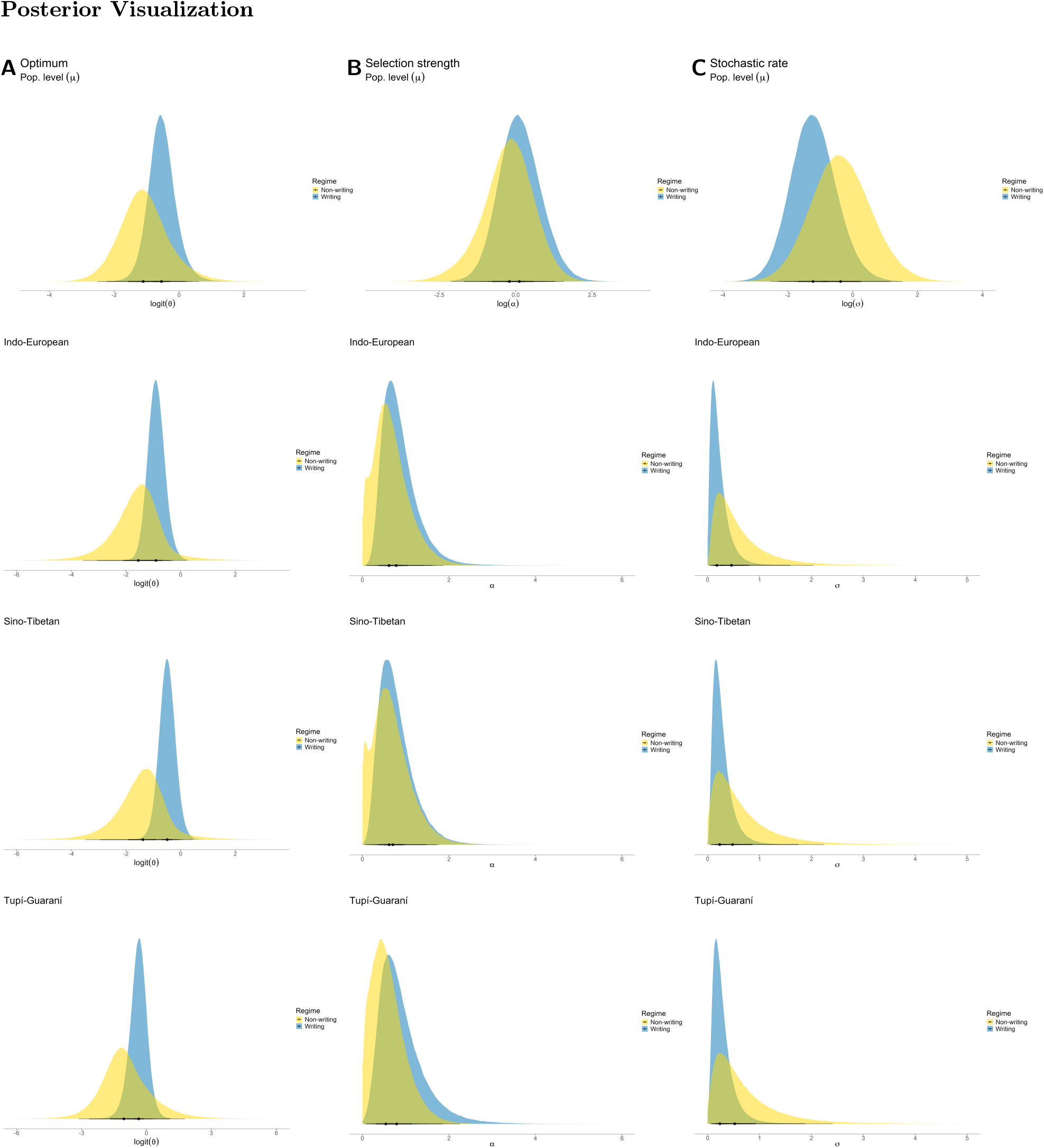
Posterior distributions of the parameters in the Ornstein–Uhlenbeck process. **A:** Posterior distributions of the optimum value (*θ*), shown for the population level (first panel), Indo-European (second), Sino-Tibetan (third), and Tupí-Guaraní (fourth). **B:** Posterior distributions of the selection strength (*α*). **C:** or distributions of the stochastic rate (*σ*).

#### Differences between the Ornstein-Uhlenbeck Regimes

**Table S5.** Differences in Ornstein-Uhlenbeck process parameter values between writing and non-writing regimes. All values in the table represent the difference between the writing and non-writing regimes, calculated as *β*_writing_ − *β*_*¬*writing_. The subscript *µ* in the *Parameter* column refers to population-level parameters, as opposed to family-level parameters (IE: Indo-European, ST: Sino-Tibetan, TG: Tupí-Guaraní). The final column, Pr(Δ*β >* 0), indicates the proportion of the posterior distribution of differences that is greater than zero – i.e., the proportion of cases in which the parameter value is higher for the writing regime than for the non-writing regime. Values above 50% indicate that the writing regime typically has higher parameter estimates (as observed for *θ* and *α*), while values below 0.5 indicate higher estimates for the non-writing regime (as seen for *σ*).

| Parameter | Median | HPDI (95%) | $\Pr(\Delta\beta) > 0$ |
| --- | --- | --- | --- |
| $\theta_\mu$ | 0.58 | $[-1.19, 2.18]$ | 76.62% |
| $\theta_{\text{IE}}$ | 0.66 | $[-0.95, 2.78]$ | 84.46% |
| $\theta_{\text{ST}}$ | 0.90 | $[-0.90, 3.08]$ | 89.45% |
| $\theta_{\text{TG}}$ | 0.69 | $[-2.13, 2.91]$ | 73.08% |
| $\alpha_\mu$ | 0.30 | $[-0.93, 1.92]$ | 74.44% |
| $\alpha_{\text{IE}}$ | 0.14 | $[-0.06, 0.41]$ | 95.42% |
| $\alpha_{\text{ST}}$ | 0.04 | $[-0.24, 0.56]$ | 63.12% |
| $\alpha_{\text{TG}}$ | 0.21 | $[-0.50, 1.28]$ | 77.34% |
| $\sigma_\mu$ | -0.84 | $[-3.14, 1.37]$ | 23.02% |
| $\sigma_{\text{IE}}$ | -0.25 | $[-1.62, 0.47]$ | 20.29% |
| $\sigma_{\text{ST}}$ | -0.22 | $[-1.72, 0.58]$ | 26.75% |
| $\sigma_{\text{TG}}$ | -0.26 | $[-1.90, 0.62]$ | 25.47% |

### Posterior Predictive Checks

**Figure S8.**
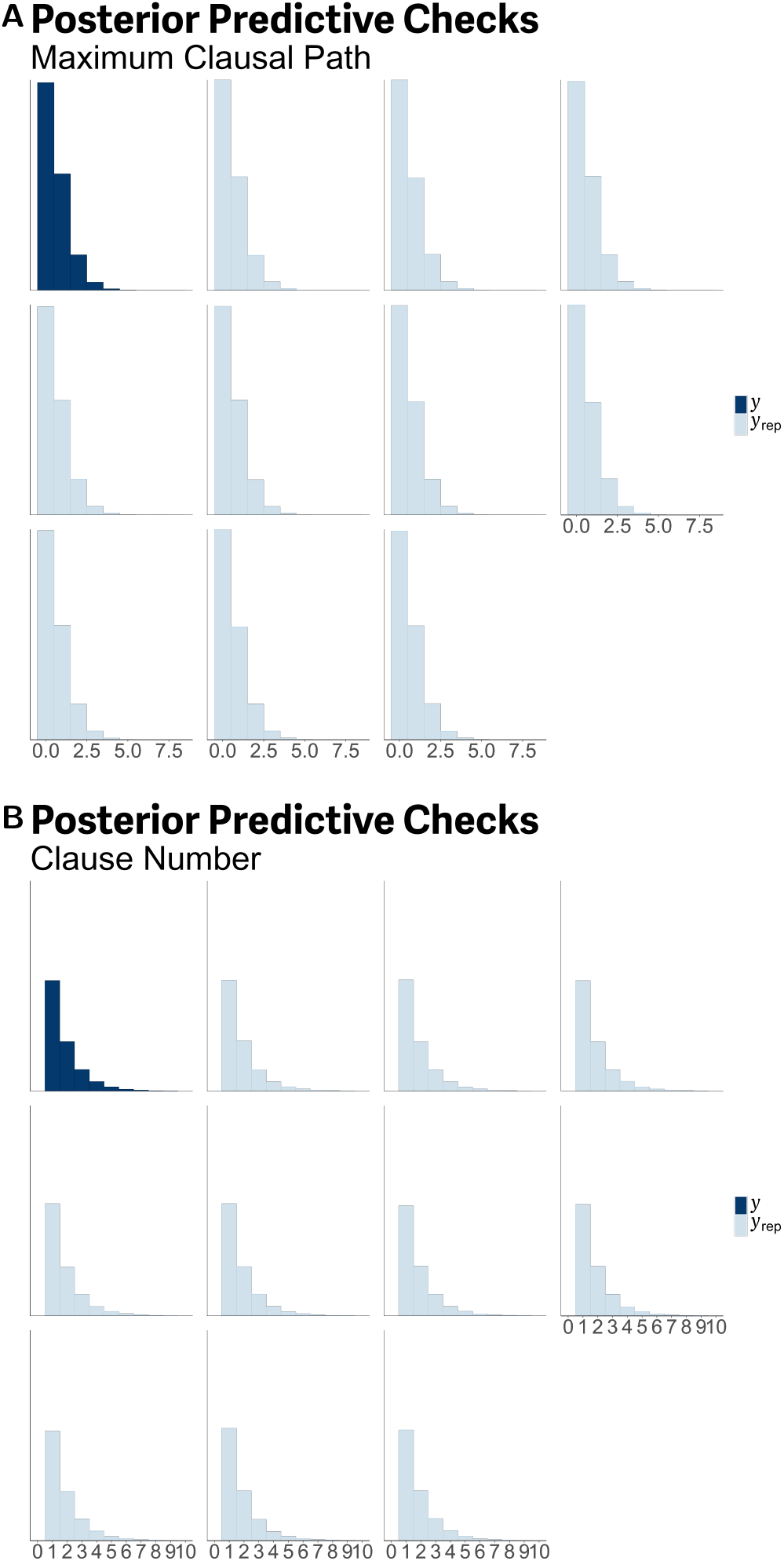
Posterior predictive checks for the phylogenetic regression models. **A:** Posterior predictive check for the clause number model. **B:** Posterior predictive check for the maximum clausal path model.

## Notes

https://doi.org/10.5281/zenodo.16107637

### Competing Interest Statement

The authors have declared no competing interest.

